# Transcriptome Analysis of Desquamative Interstitial Pneumonia (DIP)

**DOI:** 10.1101/791350

**Authors:** Shigeki Saito, Jay K. Kolls, Yaozhong Liu, Yusuke Higashi, Shinya Ohkochi, Yasuhiro Kondoh, Joseph A. Lasky, Takuji Suzuki

**Affiliations:** Department of Medicine, Tulane University School of Medicine, New Orleans, LA, USA; Department of Pediatrics, Tulane University School of Medicine, New Orleans, LA, USA; Center for Translational Research in Infection and Inflammation, Tulane University School of Medicine, New Orleans, LA, USA; Department of Biostatistics and Data Science, Tulane University School of Public Health and Tropical Medicine, New Orleans, LA, USA; Department of Medicine, Tohoku University School of Medicine, Sendai, Miyagi, Japan; Department of Medicine, Tosei General Hospital, Seto, Aichi, Japan; Department of Medicine, Jichi University School of Medicine, Shimotsuke, Tochigi, Japan

**Keywords:** DIP, RB-ILD, Alveolar Macrophage, GM-CSF, IPA, xCell

## Abstract

Desquamative interstitial pneumonia (DIP) is a rare diffuse parenchymal lung disease of unclear etiology. A recent study showed that mice overexpressing granulocyte macrophage colony-stimulating factor (GM-CSF) in lungs develop DIP-like disease, suggesting that pulmonary GM-CSF may be involved in the pathogenesis of DIP. To determine if GM-CSF is involved in human DIP lungs, we performed transcriptome analysis on human DIP lung tissue. We also extended transcriptome analysis to respiratory bronchiolitis-associated interstitial lung disease (RB-ILD), which has been thought to be in the same spectrum of disease. The analysis revealed that DIP has a distinct transcriptome profile compared to both RB-ILD and non-diseased lung controls. It also suggested that GM-CSF was a key upstream regulator in the DIP transcriptome and that the GM-CSF signaling pathway was highly activated in DIP tissue. Further bioinformatics analysis using xCell, a novel computational method that assesses enrichments of individual cell types based on gene expression, suggested that DIP is enriched for gene signatures of macrophages and other immune cells such as dendritic cells and B cells. In conclusion, our analysis shows that DIP is characterized by a GM-CSF signature, and thus GM-CSF is likely to be involved in the pathogenesis of DIP. Our analysis also suggests that immune cells other than alveolar macrophages, such as B cells, may also be involved in the pathogenesis of DIP.

## INTRODUCTION

Desquamative interstitial pneumonia (DIP) is a rare diffuse parenchymal lung disease characterized by marked accumulation of alveolar macrophages (AMs) and emphysema without extensive fibrosis or neutrophilic inflammation (1). DIP typically occurs in smokers, although it can occasionally occur in nonsmokers (2). Recently, SPC-CSF2 mice, which overexpress granulocyte macrophage colony-stimulating factor (GM-CSF, encoded by CSF2 gene) under the surfactant protein C (SPC) promoter, have been shown to display cardinal features of DIP, including AM accumulation, emphysema, secondary polycythemia, and increased mortality. These results suggest that pulmonary GM-CSF may be involved in the pathogenesis of DIP. A proposed mechanism is as follows: smoke inhalation (or another initiator) → pulmonary GM-CSF hypersecretion response → AM accumulation and activation (e.g., STAT5 phosphorylation) → MMP secretion (e.g., MMP9, MMP12) by AMs → parenchymal lung damage (i.e., DIP, emphysema) (1). To determine whether GM-CSF signaling is truly involved in human DIP lungs, we performed transcriptome analysis of human DIP lungs and controls. We also extended transcriptome analysis to respiratory bronchiolitis-associated interstitial lung disease (RB-ILD), which has been thought to be in the same spectrum of smoking-related pulmonary disease (3). Our analysis reveals that the transcriptome of DIP lungs is distinct from that of RB-ILD and controls. It also supports the concept that abnormally increased pulmonary GM-CSF signaling may play a key role in the pathogenesis of DIP. Furthermore, it suggests that immune cells other than alveolar macrophages, such as B cells, may be involved in the pathogenesis of DIP.

## METHODS

### Data source and analysis of differentially expressed genes

Microarray data of 4 DIP, 11 RB-ILD, and 50 controls in GSE32537 in the NCBI Gene Expression Omnibus (GEO) repository were used for this analysis. Yang et al. obtained DIP and RB-ILD lung tissue from the Lung Tissue Research Consortium (LTRC, a biobank created by the National Heart Lung and Blood Institute [NHLBI]), and non-diseased lung tissue (control) from the International Institute for Advancement of Medicine. The control individuals had suffered brain death, failed selection criteria for transplantation, and had no evidence of acute or chronic lung disease. For both the diseased lung tissue (i.e., DIP, RB-ILD) and the non-diseased lung tissue (i.e., control), total RNA was isolated and mRNA microarray was performed using the Human Gene 1.0 ST Array (Affymetrix) platform, and the microarray data was uploaded to GEO (4). To identify genes (transcripts) that are differentially expressed between DIP and control lung tissue, the expression data were analyzed using limma (Linear Models for Microarray Analysis) R package implemented in GEO2R (5). The Benjamini–Hochberg false discovery rate (FDR) method was used for multiple-testing corrections.

### Ingenuity Pathway Analysis (IPA) to identify canonical pathways and upstream regulators

Pathway analysis of the gene list was carried out using Ingenuity Pathway Analysis (IPA) (6). Using its extensive database of published data as well as natural language processing and curated text mining of the published literature, IPA identifies pathways likely to be responsible for observed difference in gene expression profiles. IPA can also identify putative molecular drivers (upstream regulators) of differentially expressed genes. For analysis of both canonical pathways and upstream regulators, absolute z-score ≥ 2.0 corresponds to significant changes in activity; z-score ≥ 2 implies significant activation, whereas z-score ≤ −2 implies significant inhibition. The predicted upstream regulators may themselves be differentially expressed, though this is not a criterion for inclusion (7).

### Estimation of immune cell landscape

To characterize the immune cell landscape in DIP, enrichment of various types of immune cells in DIP, RB-ILD, and controls were estimated using xCell (8), a novel computational method that assesses enrichment of individual cell types based on gene expression profile. An xCell matrix comprising gene expression profiles of 64 cell types was used as the reference matrix. Among the 64 cell types, immune cells, platelets, and erythrocytes present in lungs were selected for further analysis. Enrichment scores for each cell type in DIP, RB-ILD, and the control were compared using Kruskal-Wallis test followed by Dunn’s multiple comparison. A p-value <0.05 was considered statistically significant. Statistical analysis was performed using GraphPad Prism 7 software.

## RESULTS

### Demographic characteristics

Table 1 summarizes demographic and clinical characteristics of the DIP patients, RB-ILD patients, and the non-diseased control cohort. There are no statistically significant differences in sex or smoking status between the groups. Although there are no statistically significant differences, there is trend that DIP patients and RB-ILD patients smoked more cigarettes than controls, as expected. There is also a trend indicating that DIP patients had worse lung function than RB-ILD patients, as expected.

**Table 1.** Demographics.

| **Disease group** | **Control** | **DIP** | **p value (vs Control)** | **RB-ILD** | **p value (vs Control)** | **p value (vs DIP)** |
| --- | --- | --- | --- | --- | --- | --- |
| Number | 50 | 4 |  | 11 |  |  |
| Age: mean (SD) | 45.6 (18.6) | 50 (7.0) | 0.83* | 52.2 (11.0) | 0.35* | 0.65* |
| Sex: N (%) | M 27 (54%)<br>F 23 (46%) | M 3 (75%),<br>F 1 (25%) | 0.62† | M 4 (36%),<br>F 7 (64%) | 0.34† | 0.28† |
| Smoking status: N (%) |  |  | 0.64†† |  | 1.0†† | 1.0†† |
| -Current | 21 (42%) | 1 (25%) |  | 1 (9%) |  |  |
| -Former | 7 (14%) | 2 (50%) |  | 6 (55%) |  |  |
| -Never | 20 (40%) | 1 (25%) |  | 4 (36%) |  |  |
| -Unknown | 2 (4%) | 0 (0%) |  | 0 (0%) |  |  |
| Pack-year: mean (SD) | 12.0 (18.1)<br>(6 missing values) | 24.0 (17.1) | 0.12* | 33.9 (35.9) | 0.08* | 0.95* |
| FVC: mean (SD) | Unknown | 67.4 (8.5) | N/A | 85.7 (13.1)<br>(1 missing values) | N/A | 0.07 |
| DLCO: mean (SD) | unknown | 54.3 (0)<br>(3 missing values) | N/A | 77.1 (21.3)<br>(2 missing values) | N/A |  |

SD: standard deviation, M: male, F: female, N: Number, FVC: forced vital capacity, DLCO: diffusing capacity of the lungs for carbon monoxide
\*By Mann-Whitney test.
†By Fischer's exact test
††By Fischer's exact test for Current + Former vs Never

### Canonical pathways and upstream regulators responsible for differentially expressed transcripts

#### DIP to control comparison

There were 1932 transcript IDs meeting FDR <0.05 cut-off, and 438 transcript IDs meeting FDR <0.05 and absolute fold change ≥□2 cut-off (207 upregulated, 231 downregulated). IPA identified multiple canonical pathways that were significantly overrepresented in DIP compared to control (Table 2). Many of them were related to innate or adaptive immunity. For example, pathways related to innate immunity include “Granulocyte Adhesion and Diapedesis,” “Leukocyte Extravasation Signaling” (with significant z-score: 2.714), “Dendritic Cell Maturation,” “Phagosome Formation,” “Phagosome Maturation,” and “Complement System.” Pathways related to adaptive immunity include “Th1 and Th2 Activation Pathway,” “Th1 Pathway,” “Th2 Pathway,” “T Helper Cell Differentiation,” “T Cell Exhaustion Signaling Pathway,” and “B Cell Development.” Pathways related to both innate and adaptive immunity included “Antigen Presentation Pathway,” “Communication between Innate and Adaptive Immune Cells,” and “PD-1, PD-L1 cancer immunotherapy pathway.” Of note, the “Hepatic Fibrosis/Hepatic Stellate Cell Activation” pathway was found to be significantly deregulated. This appears consistent with the fact that a varying degree of fibrosis is observed in DIP. Furthermore, “Inhibition of Matrix Metalloproteinases (MMPs)” (i.e., counteraction by TIMPs) was found to be significantly inhibited (Z-score-2.449), which suggests that MMPs are significantly activated (Supplemental Figure 1). This supports the concept that MMPs may be involved in the pathogenesis of DIP (1).

**Table 2.** Canonical Pathways in DIP (compared to Control)

| **Ingenuity Canonical Pathways** | **-log(p-value)** | **z-score** |
| --- | --- | --- |
| Granulocyte Adhesion and Diapedesis | 1.04E+01 |  |
| Hepatic Fibrosis / Hepatic Stellate Cell Activation | 5.56E+00 |  |
| Primary Immunodeficiency Signaling | 5.22E+00 |  |
| Inhibition of Matrix Metalloproteases | 5.14E+00 | -2.449 |
| Osteoarthritis Pathway | 4.95E+00 | -0.258 |
| LPS/IL-1 Mediated Inhibition of RXR Function | 4.93E+00 | -1.134 |
| Th1 and Th2 Activation Pathway | 4.83E+00 |  |
| Role of Osteoblasts, Osteoclasts and Chondrocytes in Rheumatoid Arthritis | 4.66E+00 |  |
| Agranulocyte Adhesion and Diapedesis | 4.52E+00 |  |
| Th2 Pathway | 4.46E+00 | 1.633 |
| Allograft Rejection Signaling | 4.36E+00 |  |
| STAT3 Pathway | 4.18E+00 | -1.89 |
| Communication between Innate and Adaptive Immune Cells | 4.12E+00 |  |
| Leukocyte Extravasation Signaling | 4.00E+00 | 2.714 |
| Oncostatin M Signaling | 3.77E+00 | 0.447 |
| Phagosome Formation | 3.51E+00 |  |
| Atherosclerosis Signaling | 3.28E+00 |  |
| Altered T Cell and B Cell Signaling in Rheumatoid Arthritis | 3.10E+00 |  |
| Dendritic Cell Maturation | 2.92E+00 | 3 |
| Th1 Pathway | 2.87E+00 | 0 |
| LXR/RXR Activation | 2.63E+00 | 1.89 |
| Hematopoiesis from Pluripotent Stem Cells | 2.54E+00 |  |
| Autoimmune Thyroid Disease Signaling | 2.49E+00 |  |
| Tryptophan Degradation to 2-amino-3-carboxymuconate Semialdehyde | 2.27E+00 |  |
| Role of Macrophages, Fibroblasts and Endothelial Cells in Rheumatoid Arthritis | 2.24E+00 |  |
| NF-κB Signaling | 2.19E+00 | -1.667 |
| SPINK1 General Cancer Pathway | 2.15E+00 | 2.236 |
| Hepatic Cholestasis | 2.12E+00 |  |
| OX40 Signaling Pathway | 2.02E+00 |  |
| B Cell Development | 2.02E+00 |  |
| IL-10 Signaling | 1.99E+00 |  |
| Bladder Cancer Signaling | 1.98E+00 |  |
| FXR/RXR Activation | 1.97E+00 |  |
| PTEN Signaling | 1.95E+00 | 2.646 |
| PPAR Signaling | 1.88E+00 | 0.816 |
| Iron homeostasis signaling pathway | 1.87E+00 |  |
| Nicotine Degradation II | 1.86E+00 | -2 |
| IL-8 Signaling | 1.84E+00 | -0.333 |
| Antigen Presentation Pathway | 1.70E+00 |  |
| NAD biosynthesis II (from tryptophan) | 1.66E+00 |  |
| Inhibition of Angiogenesis by TSP1 | 1.63E+00 |  |
| Bile Acid Biosynthesis, Neutral Pathway | 1.60E+00 |  |
| Chondroitin Sulfate Degradation (Metazoa) | 1.53E+00 |  |
| cAMP-mediated signaling | 1.53E+00 | -1.89 |
| Graft-versus-Host Disease Signaling | 1.52E+00 |  |
| Systemic Lupus Erythematosus Signaling | 1.51E+00 |  |
| T Cell Exhaustion Signaling Pathway | 1.50E+00 | -1 |
| Complement System | 1.49E+00 |  |
| Dermatan Sulfate Degradation (Metazoa) | 1.48E+00 |  |
| T Helper Cell Differentiation | 1.45E+00 |  |
| PD-1, PD-L1 cancer immunotherapy pathway | 1.44E+00 |  |
| Sulfate Activation for Sulfonation | 1.42E+00 |  |
| Cysteine Biosynthesis/Homocysteine Degradation | 1.42E+00 |  |
| Dermatan Sulfate Biosynthesis (Late Stages) | 1.37E+00 |  |
| Caveolar-mediated Endocytosis Signaling | 1.37E+00 |  |
| Phagosome Maturation | 1.34E+00 |  |
| TREM1 Signaling | 1.33E+00 | 1 |

IPA also predicted many upstream regulators responsible for transcriptome changes in DIP. In the “cytokines” category, four genes were predicted to be upstream regulators with a significant Z-score: *CSF2*, *IL4*, *IL5*, and *SPP1* (osteopontin) (Table 3). Among these, *CSF2* (the gene encoding GM-CSF) had the highest activation Z-score of 3.162 (i.e., predicted to be significantly activated in DIP compared to control), although its expression was not found to be upregulated in DIP (Figures 1, 2). This supports the notion that GM-CSF may be involved in the pathogenesis of DIP (1). *SPP1* also had a high activation Z-score (2.335), and its expression was upregulated in DIP (log expression ratio [DIP vs. control] = 2.878). This is consistent with the previous study, which showed increased osteopontin expression in, and secretion from, AMs in DIP lungs (9).

**Figure 1.**
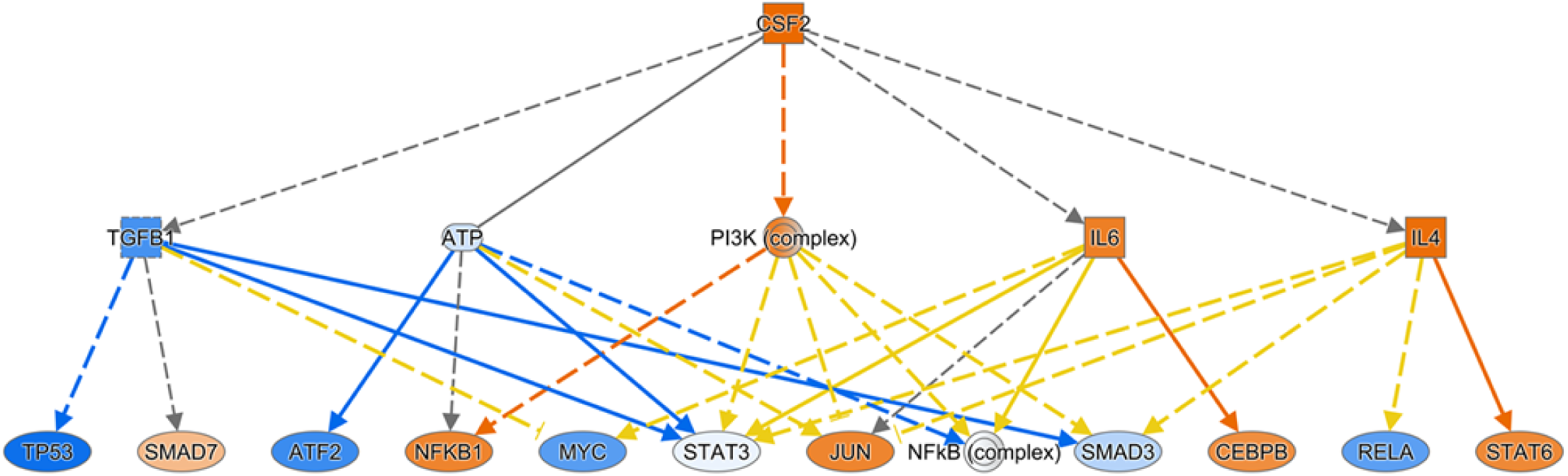
*CSF2* is a predicted upstream regulator driving pathway activation in DIP transcriptome. *CSF2* was predicted by IPA as an upstream regulator driving pathway activation (i.e., *CSF2* was predicted to be activated) (activation z-score = +3.162). The *CSF2* expression level itself was not significantly different between DIP and control. The genes/regulators are colored according to their predicted activation state: activated (orange) or inhibited (blue). Darker colors indicate higher absolute Z-scores. The edges connecting the nodes are colored orange when leading to activation of the downstream node, blue when leading to its inhibition, yellow if the findings underlying the relationship are inconsistent with the state of the downstream node, and grey if the effect is not predicted. Pointed arrowheads indicate that the downstream node is expected to be activated if the upstream node connected to it is activated, while blunt arrowheads indicate that the downstream node is expected to be inhibited if the upstream node that connects to it is activated. The dashed lines indicate virtual relationships composed of the net effect of the paths between the root regulator and the target genes (6).

**Figure 2.**
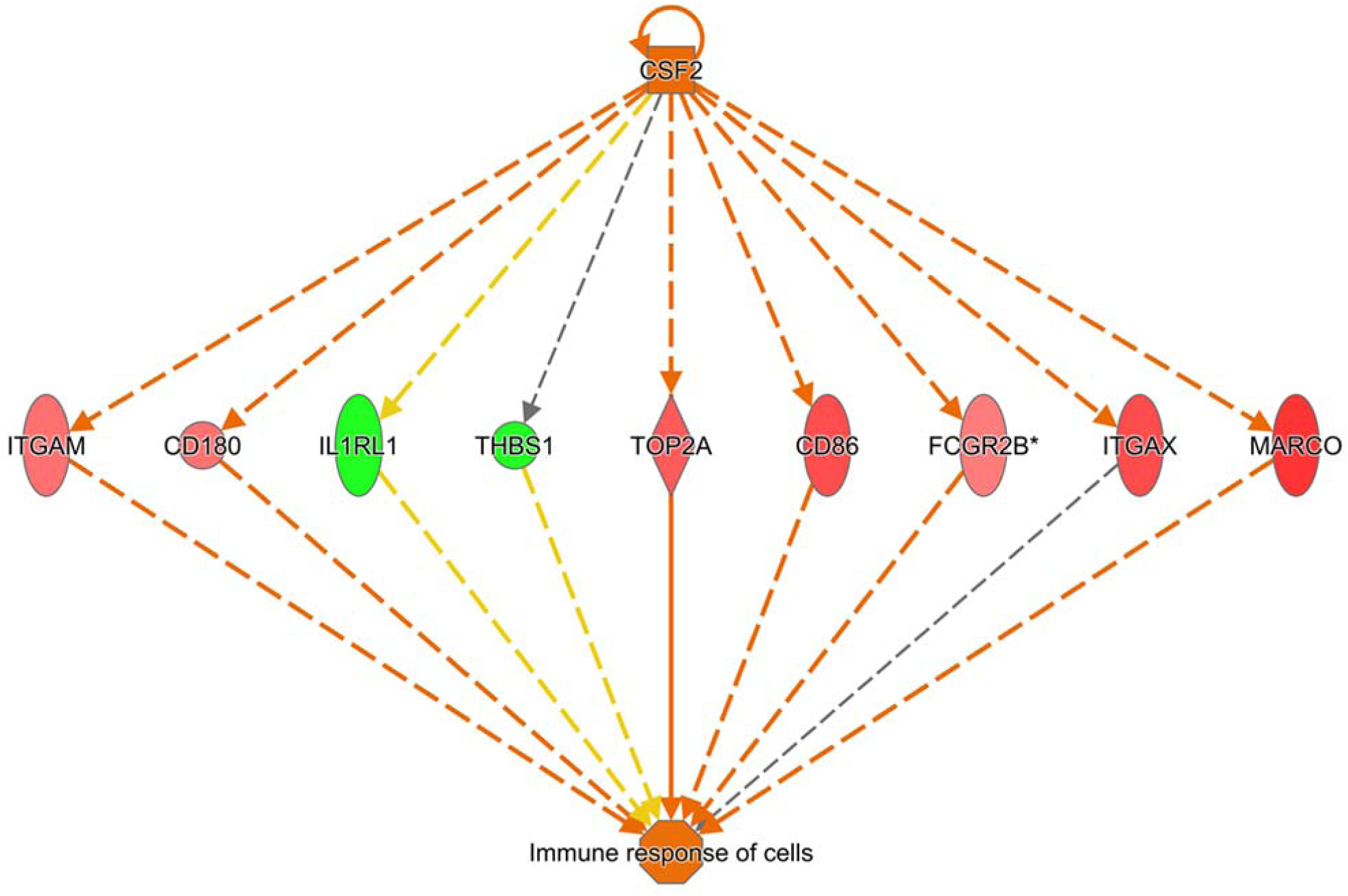
Predicted regulatory effects of *CSF2*. Upregulated (red) or downregulated (green) genes are indicated in the network surrounding *CSF2*. Arrows are colored as in Figure 1.

In the “transcription regulators” category, 13 transcription factors were identified as potential upstream regulators: Five were predicted to be significantly inhibited (*ZBTB17, ZFP36, EGR2, NUPR1, NANOG*), and eight were predicted to be significantly activated (*NEUROG1, ETV5, CLOCK, KEAP1, SOX3, WT1, PPARGC1A, RUNX3*). In the “ligand-dependent nuclear factor” category, *NR3C1* (gene encoding glucocorticoid receptor) was predicted to be significantly inhibited (Z-score −2.235) (Figure 3). This may be reflecting the fact that corticosteroids are used as an effective treatment in some cases of DIP (10). In contrast, *PPARA* was predicted to be significantly activated (Z-score 2.159).

**Figure 3.**
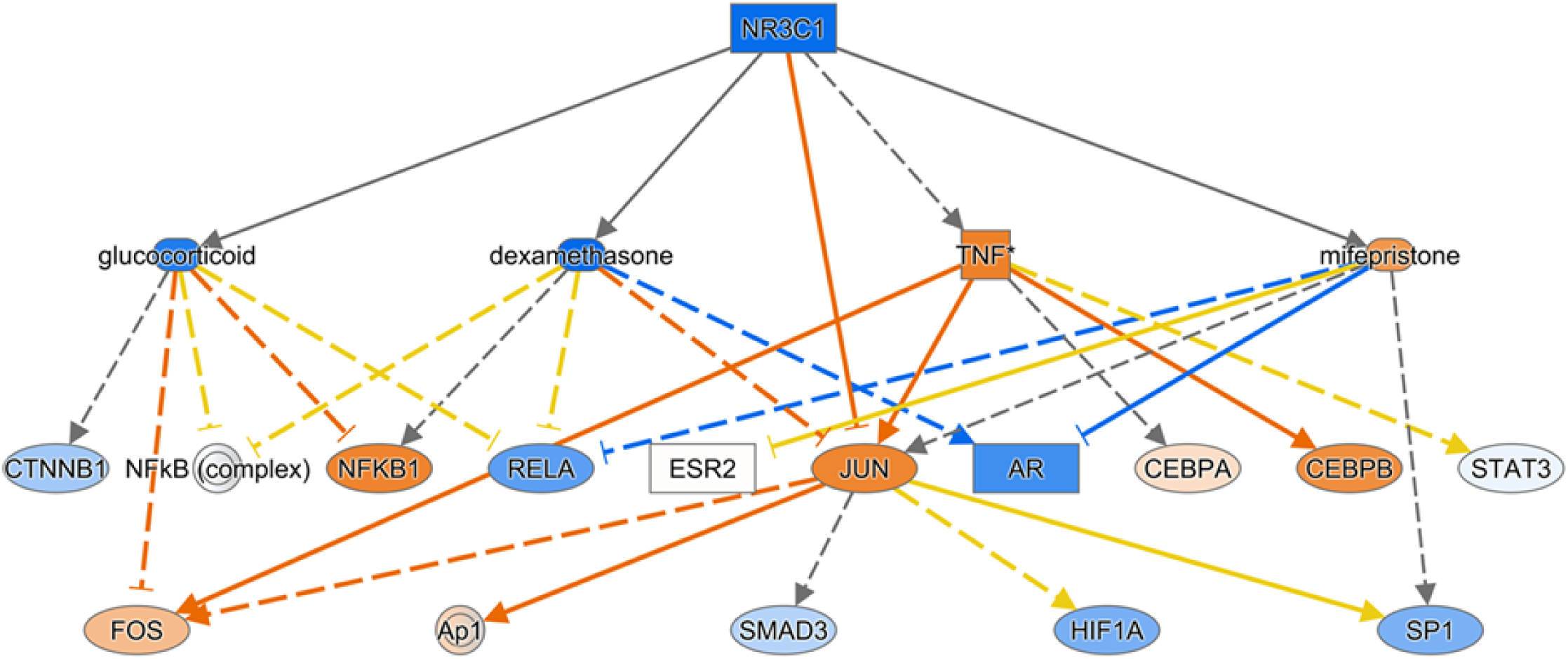
*NR3C1* (corticosteroid receptor) is predicted to be one of the inhibited upstream regulators in the DIP transcriptome. Genes/regulators and arrows are colored as in Figure 1.

In the “transmembrane receptors” category, *CD40* (expressed on antigen presenting cells, including dendritic cells, macrophages, and B cells; Supplemental Figure 2), *TLR2*, and *TREM1* (expressed on various cells of the myeloid lineage) were identified as potential upstream regulators. All were predicted to be significantly activated.

In the “complex” category, BCR was the only upstream regulator with a significant Z-score of 2.174 (i.e., predicted to be activated) (Supplemental Figure 3).

IPA also identified some drugs and chemicals as potential upstream regulators. For example, glucocorticoids such as dexamethasone (Supplemental Figure 4), fluticasone propionate, triamcinolone acetonide, and fluocinolone acetonide were predicted to be significantly inhibited in DIP compared to control. This supports the fact that corticosteroids are an effective treatment in some cases of DIP. Interestingly, filgrastim (granulocyte-colony stimulating factor), chloroquine (Supplemental Figure 5), 4-hydroxytamoxifen (a selective estrogen receptor modulator), Z-LLL-CHO (also known as MG-132, a proteasome inhibitor), and lactacystin (a proteasome inhibitor), were predicted to be significantly inhibited. In contrast, diethylstilbestrol (DES; a synthetic form of estrogen), enterolactone (a phytoestrogen), daidzein (a phytoestrogen), chrysotile asbestos, cephaloridine (cephalosporin), pioglitazone (an agonist for PPAR-γ [and PPAR-α, to lesser degree]), and cholesterol were predicted to be significantly activated.

#### RB-ILD to control comparison

There were 1767 transcript IDs meeting FDR <0.05 cut-off, and 100 transcript IDs meeting FDR <0.05 and absolute fold change ≥□2 cut-off (31 upregulated, 69 downregulated).

IPA identified pathways that are significantly overrepresented in RB-ILD compared to control, although the number of pathways in this comparison was less than the number of pathways in the DIP to control comparison (Table 4). Unlike the DIP vs control comparison, the RB-ILD vs control comparison did not suggest pathways directly linked to adaptive immunity (e.g., B cells, T cells). Pathways identified in both DIP and RB-ILD include “Granulocyte Adhesion and Diapedesis,” “Hepatic Fibrosis/Hepatic Stellate Cell Activation,” “Role of Osteoblasts, Osteoclasts and Chondrocytes in Rheumatoid Arthritis,” “Role of Macrophages, Fibroblasts and Endothelial Cells in Rheumatoid Arthritis,” “PPAR Signaling,” “STAT3 Pathway,” “LXR/RXR Activation,” and “LPS/IL-1 Mediated Inhibition of RXR Function.”

**Table 4.**
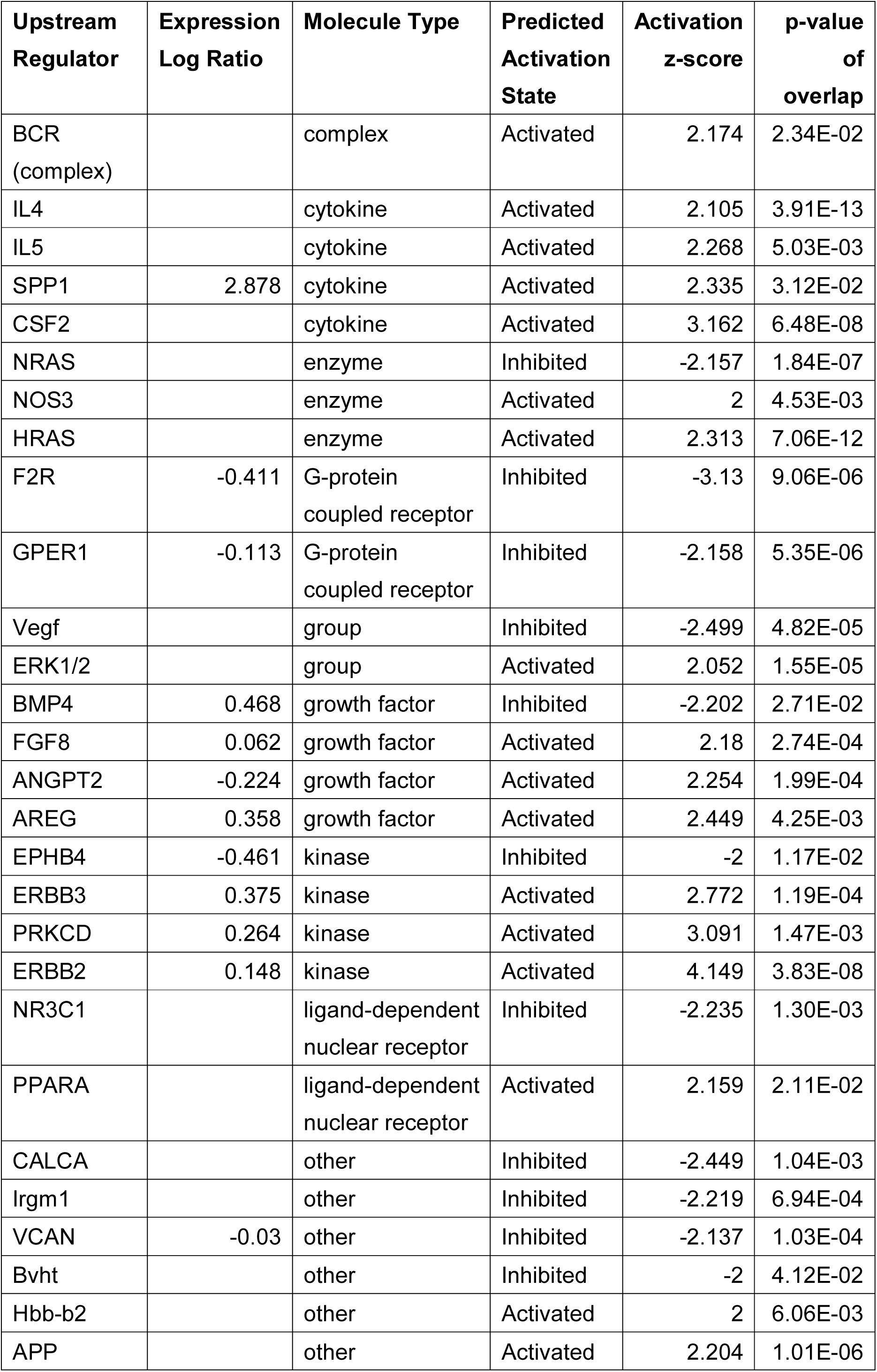
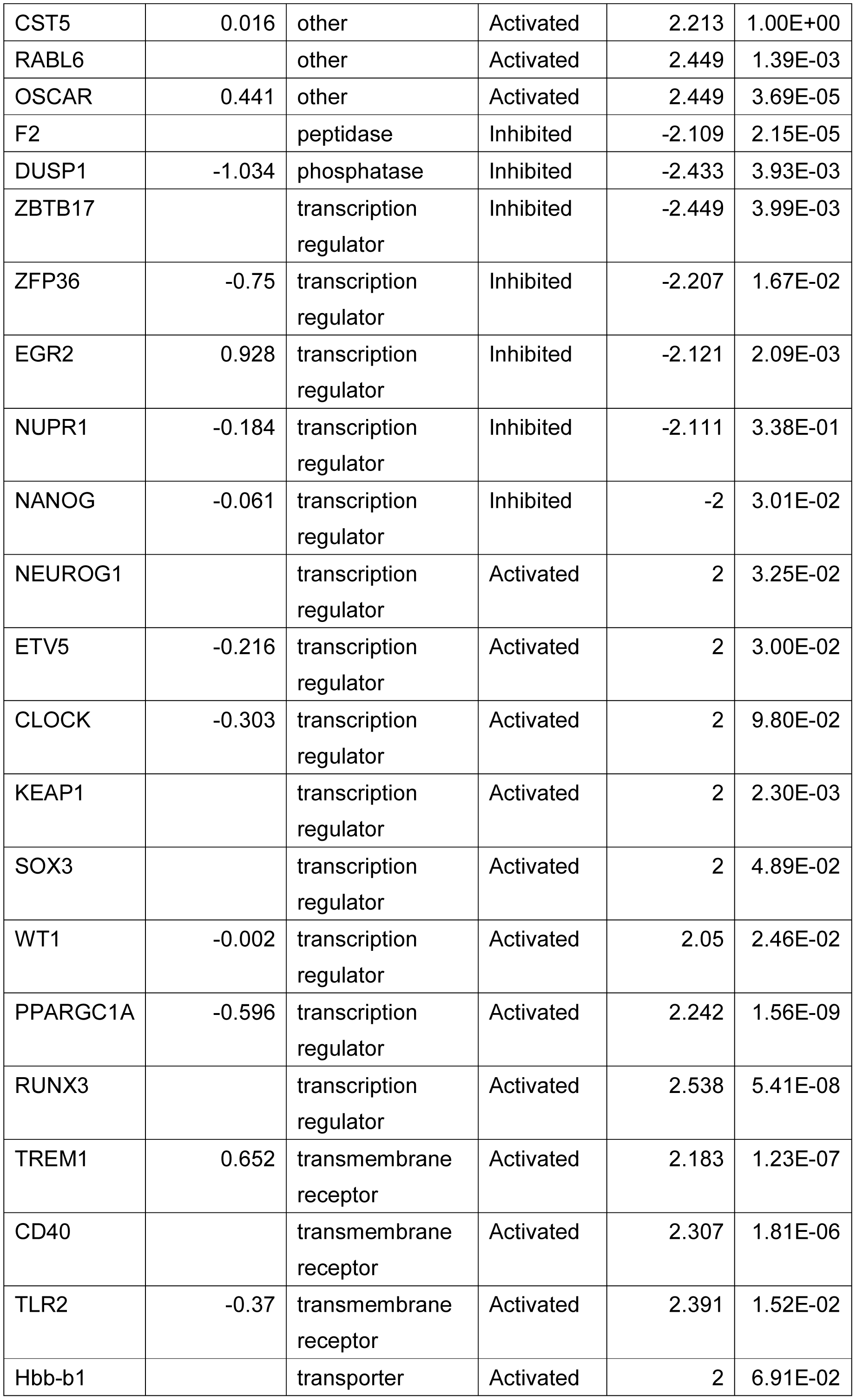
Upstream Regulators in DIP (compared to Control)

Upstream regulators identified/predicted in RB-ILD differ from the ones identified/predicted in DIP (Table 5). For example, in the “cytokines” category, the following were predicted to be upstream regulators with significant Z-scores: *TNF, IL6, IL1A, IL1B, TNFSF12, OSM, IL17A, IFNA2*. All of these were predicted to be significantly inhibited (i.e., Z-score <-2). Consistently, *STAT3* (in “transcription regulator” category) was predicted to be significantly inhibited (Z-score: −2.292). Of note, *CSF2* was not identified as an upstream regulator.

**Table 5.** Canonical Pathway in RB-ILD (compared to Control)

| **Ingenuity Canonical Pathways** | **-log(p-value)** | **z-score** |
| --- | --- | --- |
| Role of Macrophages, Fibroblasts and Endothelial Cells in Rheumatoid Arthritis | 4.01E+00 |  |
| Granulocyte Adhesion and Diapedesis | 3.91E+00 |  |
| SPINK1 General Cancer Pathway | 3.64E+00 | 2 |
| Hepatic Fibrosis / Hepatic Stellate Cell Activation | 3.60E+00 |  |
| Osteoarthritis Pathway | 3.34E+00 | -2.236 |
| STAT3 Pathway | 3.31E+00 |  |
| Role of Osteoblasts, Osteoclasts and Chondrocytes in Rheumatoid Arthritis | 3.21E+00 |  |
| PPAR Signaling | 2.86E+00 | 2 |
| Heparan Sulfate Biosynthesis (Late Stages) | 2.60E+00 |  |
| LXR/RXR Activation | 2.57E+00 | 1 |
| LPS/IL-1 Mediated Inhibition of RXR Function | 2.50E+00 |  |
| Heparan Sulfate Biosynthesis | 2.46E+00 |  |
| Iron homeostasis signaling pathway | 2.44E+00 |  |
| IL-10 Signaling | 2.36E+00 |  |
| Apelin Cardiac Fibroblast Signaling Pathway | 2.26E+00 |  |
| Cysteine Biosynthesis/Homocysteine Degradation | 2.02E+00 |  |
| Coagulation System | 1.91E+00 |  |
| Triacylglycerol Degradation | 1.84E+00 |  |
| p38 MAPK Signaling | 1.75E+00 |  |
| GP6 Signaling Pathway | 1.72E+00 |  |
| NAD Biosynthesis III | 1.72E+00 |  |
| Glucocorticoid Receptor Signaling | 1.66E+00 |  |
| IL-6 Signaling | 1.64E+00 |  |
| Serine Biosynthesis | 1.62E+00 |  |
| Lysine Degradation II | 1.62E+00 |  |
| Superpathway of Serine and Glycine Biosynthesis I | 1.48E+00 |  |

#### DIP to RB-ILD comparison

There were 656 transcript IDs meeting FDR <0.05 cut-off, and 200 transcript IDs meeting FDR <0.05 and absolute fold change ≥□2 cut-off (81 upregulated, 119 downregulated).

IPA identified multiple pathways that are significantly overrepresented in DIP compared to RB-ILD (Table 6). These included some of the pathways identified in the DIP to control comparison, such as “Granulocyte Adhesion and Diapedesis,” “Agranulocyte Adhesion and Diapedesis,” “Inhibition of Matrix Metalloproteases,” “Complement System,” “Leukocyte Extravasation Signaling,” “Phagosome Formation,” “Hepatic Fibrosis/Hepatic Stellate Cell Activation,” “Oncostatin M Signaling,” “IL-8 Signaling,” “Altered T Cell and B Cell Signaling in Rheumatoid Arthritis,” and “Caveolar-mediated Endocytosis Signaling.”

**Table 6.**
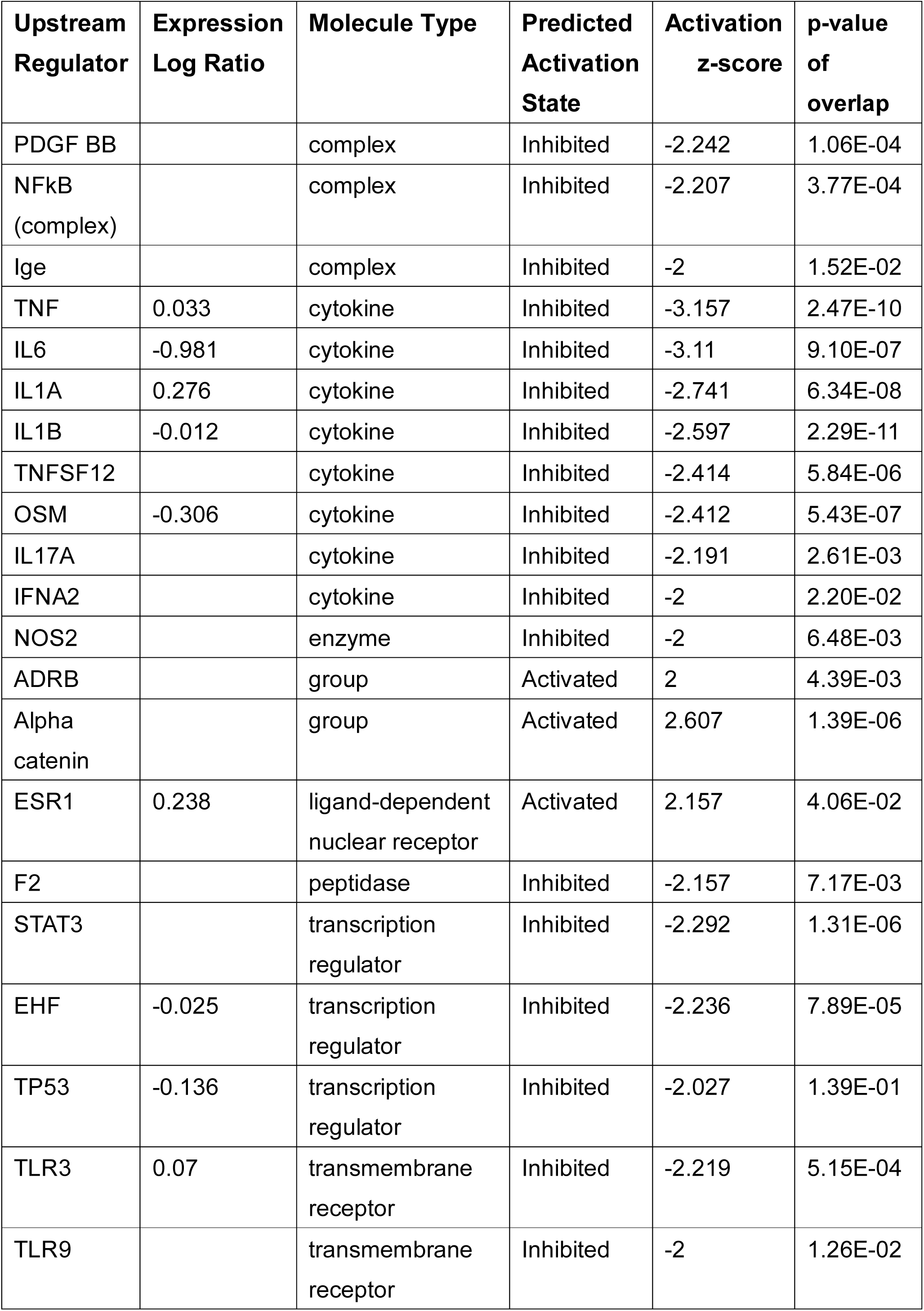
Upstream Regulators in RB-ILD (compared to Control)

| **Upstream Regulator** | **Expression Log Ratio** | **Molecule Type** | **Predicted Activation State** | **Activation z-score** | **p-value of overlap** |
| --- | --- | --- | --- | --- | --- |
| PDGF BB |  | complex | Inhibited | -2.242 | 1.06E-04 |
| NFkB (complex) |  | complex | Inhibited | -2.207 | 3.77E-04 |
| Ige |  | complex | Inhibited | -2 | 1.52E-02 |
| TNF | 0.033 | cytokine | Inhibited | -3.157 | 2.47E-10 |
| IL6 | -0.981 | cytokine | Inhibited | -3.11 | 9.10E-07 |
| IL1A | 0.276 | cytokine | Inhibited | -2.741 | 6.34E-08 |
| IL1B | -0.012 | cytokine | Inhibited | -2.597 | 2.29E-11 |
| TNFSF12 |  | cytokine | Inhibited | -2.414 | 5.84E-06 |
| OSM | -0.306 | cytokine | Inhibited | -2.412 | 5.43E-07 |
| IL17A |  | cytokine | Inhibited | -2.191 | 2.61E-03 |
| IFNA2 |  | cytokine | Inhibited | -2 | 2.20E-02 |
| NOS2 |  | enzyme | Inhibited | -2 | 6.48E-03 |
| ADRB |  | group | Activated | 2 | 4.39E-03 |
| Alpha catenin |  | group | Activated | 2.607 | 1.39E-06 |
| ESR1 | 0.238 | ligand-dependent nuclear receptor | Activated | 2.157 | 4.06E-02 |
| F2 |  | peptidase | Inhibited | -2.157 | 7.17E-03 |
| STAT3 |  | transcription regulator | Inhibited | -2.292 | 1.31E-06 |
| EHF | -0.025 | transcription regulator | Inhibited | -2.236 | 7.89E-05 |
| TP53 | -0.136 | transcription regulator | Inhibited | -2.027 | 1.39E-01 |
| TLR3 | 0.07 | transmembrane receptor | Inhibited | -2.219 | 5.15E-04 |
| TLR9 |  | transmembrane receptor | Inhibited | -2 | 1.26E-02 |

Upstream regulators identified/predicted in the DIP to RB-ILD comparison also included some of the upstream regulators identified in DIP to control comparison, such as *CSF2, IL4, SPP1*, and *NR3C1* (Table 7). Of note, *CSF2* was predicted to be strongly activated (Z-score 3.418), and *NR3C1* was predicted to be strongly inhibited (Z-score −2.213).

**Table 7.** Canonical Pathway in DIP (compared to RB-ILD)

| **Ingenuity Canonical Pathways** | **-log(p-value)** | **z-score** |
| --- | --- | --- |
| Granulocyte Adhesion and Diapedesis | 6.12E+00 |  |
| Agranulocyte Adhesion and Diapedesis | 5.83E+00 |  |
| Inhibition of Matrix Metalloproteases | 5.66E+00 | -2.449 |
| Complement System | 4.52E+00 |  |
| Axonal Guidance Signaling | 4.03E+00 |  |
| Atherosclerosis Signaling | 3.66E+00 |  |
| Osteoarthritis Pathway | 3.60E+00 | 1.667 |
| Bladder Cancer Signaling | 3.37E+00 |  |
| Leukocyte Extravasation Signaling | 3.09E+00 | 2.828 |
| Oncostatin M Signaling | 3.02E+00 | 1 |
| Phagosome Formation | 3.02E+00 |  |
| HIF1α Signaling | 3.00E+00 | |
| Hepatic Fibrosis / Hepatic Stellate Cell Activation | 2.57E+00 |  |
| Airway Pathology in Chronic Obstructive Pulmonary Disease | 2.56E+00 |  |
| IL-8 Signaling | 2.40E+00 | 1.134 |
| Caveolar-mediated Endocytosis Signaling | 2.29E+00 |  |
| Nicotine Degradation II | 1.89E+00 |  |
| Colorectal Cancer Metastasis Signaling | 1.87E+00 | 1.134 |
| Glutamate Receptor Signaling | 1.71E+00 |  |
| Reelin Signaling in Neurons | 1.34E+00 |  |
| Altered T Cell and B Cell Signaling in Rheumatoid Arthritis | 1.34E+00 |  |
| IL-17A Signaling in Fibroblasts | 1.31E+00 |  |
| dTMP De Novo Biosynthesis | 1.30E+00 |  |

**Table 8.**
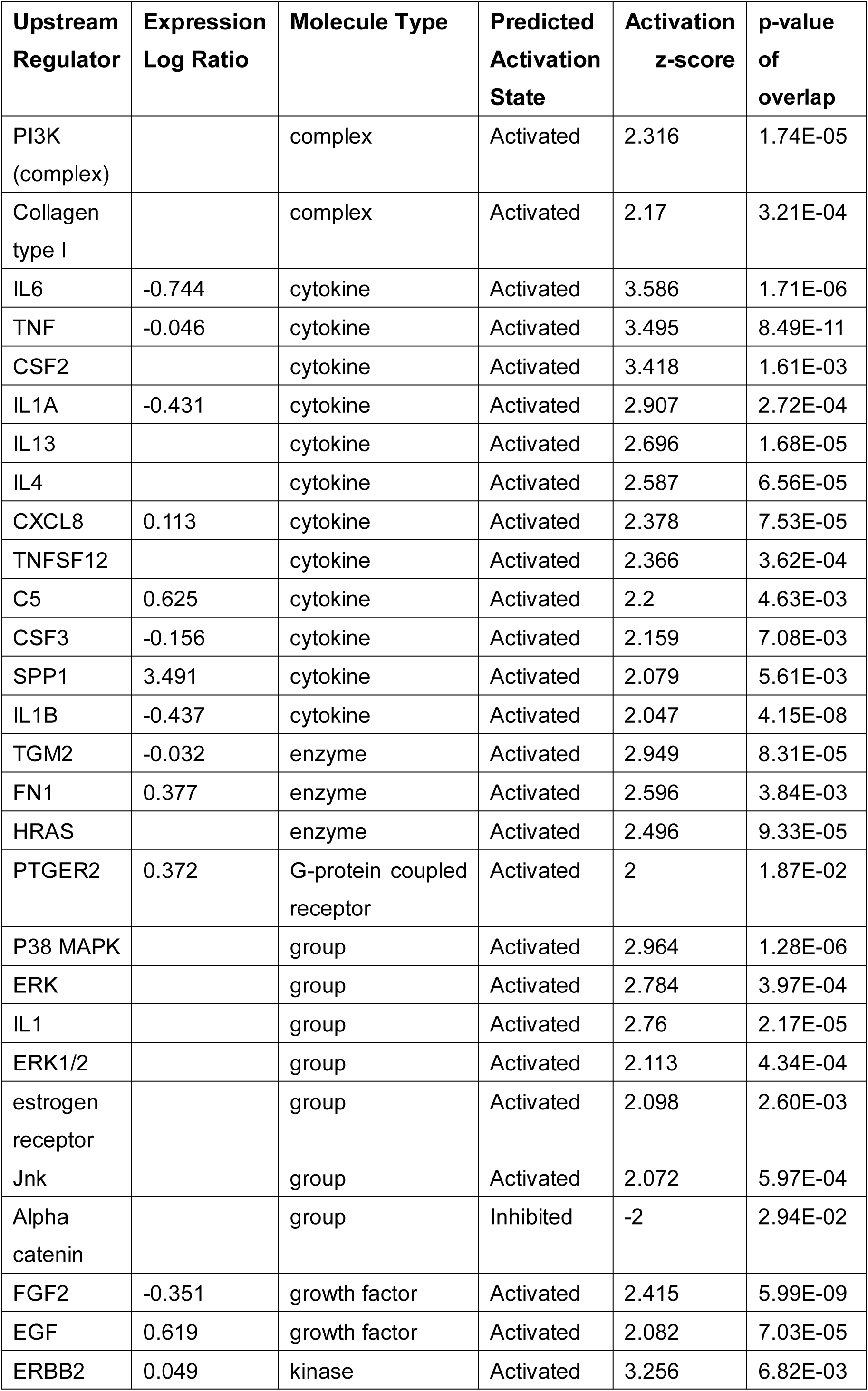
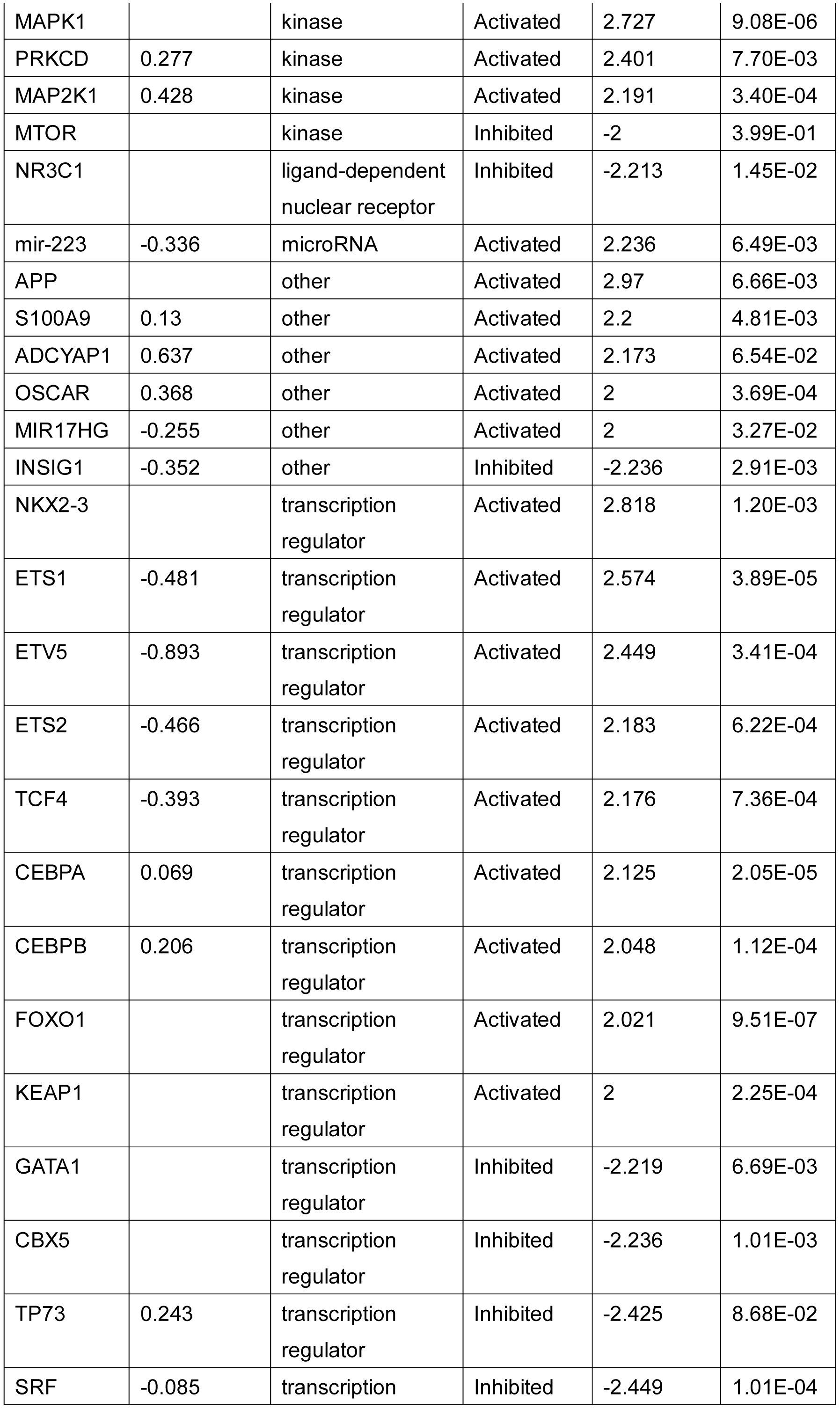
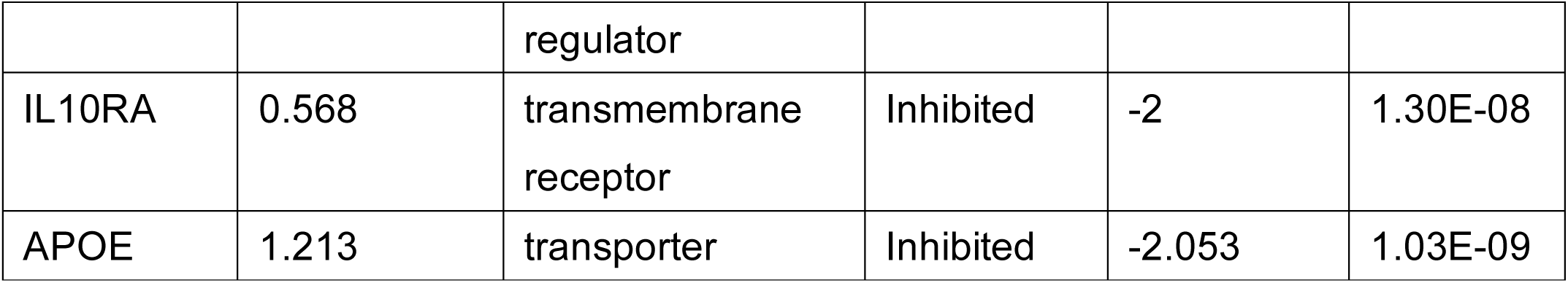
Upstream Regulators in DIP (compared to RB-ILD)

These results indicate that, in the transcriptome, both DIP and RB-ILD are distinct from control, and DIP is distinct from RB-ILD. In DIP, the GM-CSF signaling pathway and MMP pathway appear activated, as seen in SPC-CSF2 mice. Many other pathways related to innate and adaptive immunity also appear to be activated in DIP.

### Characterization of immune cell landscape in DIP lungs using xCell

#### Lymphoid cells

The gene signature of B cells was significantly enriched in DIP compared to control tissue. In particular, the gene signatures of naïve B cells, memory B cells, and class-switched B cells were significantly enriched in DIP compared to both control and RB-ILD (Figure 4). The gene signature of T cells was not significantly enriched in DIP compared to control (Figure 5), although IPA suggested overrepresentation of pathways related to T cells in DIP compared to control (Table 2). The gene signature of natural killer (NK) cells was significantly less enriched in DIP compared to RB-ILD and control (Figure 6).

**Figures 4-10.**
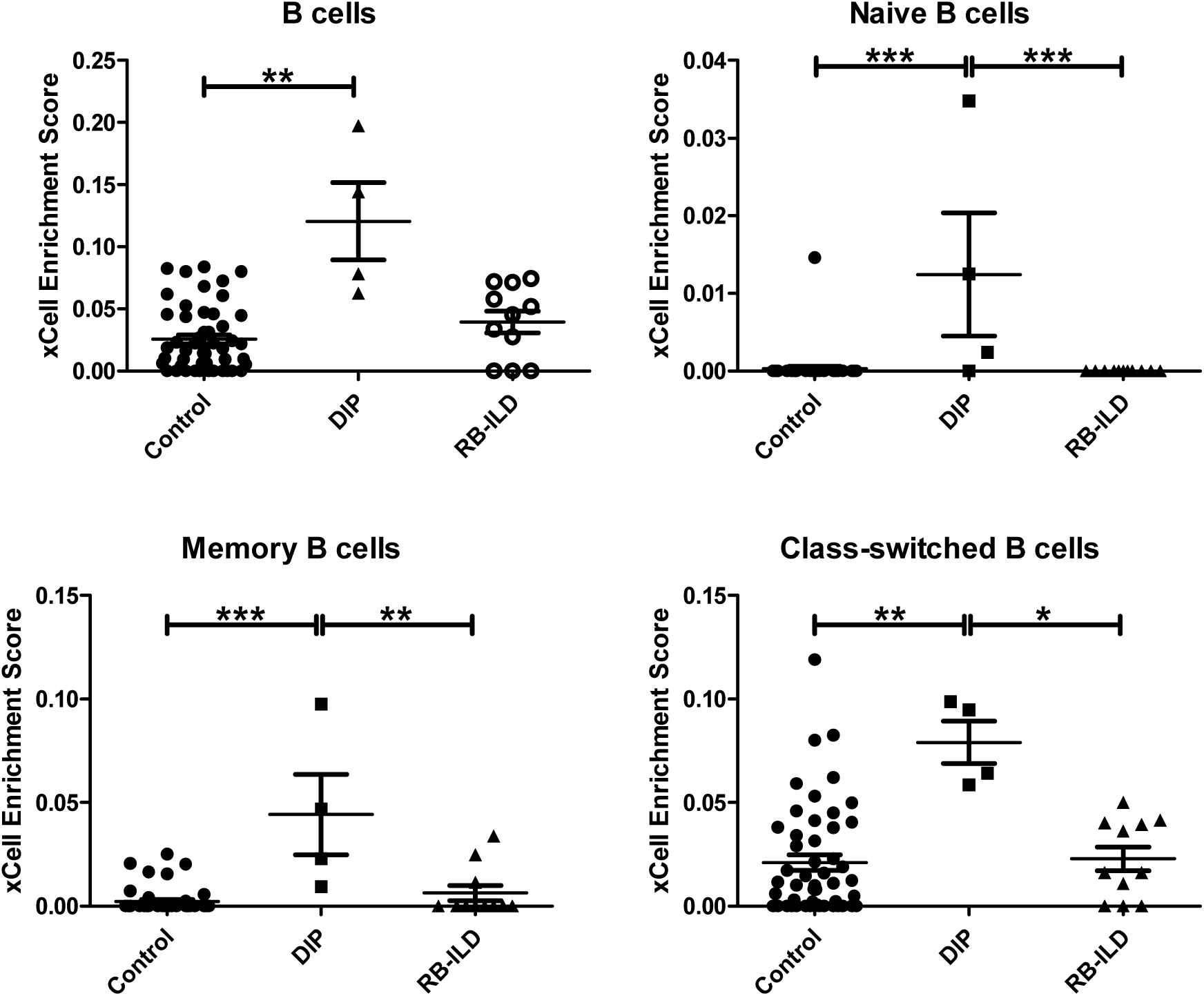
xCell-inferred enrichment scores for cell types according to the disease states. Figure 4. B cells

**Figure 5.**
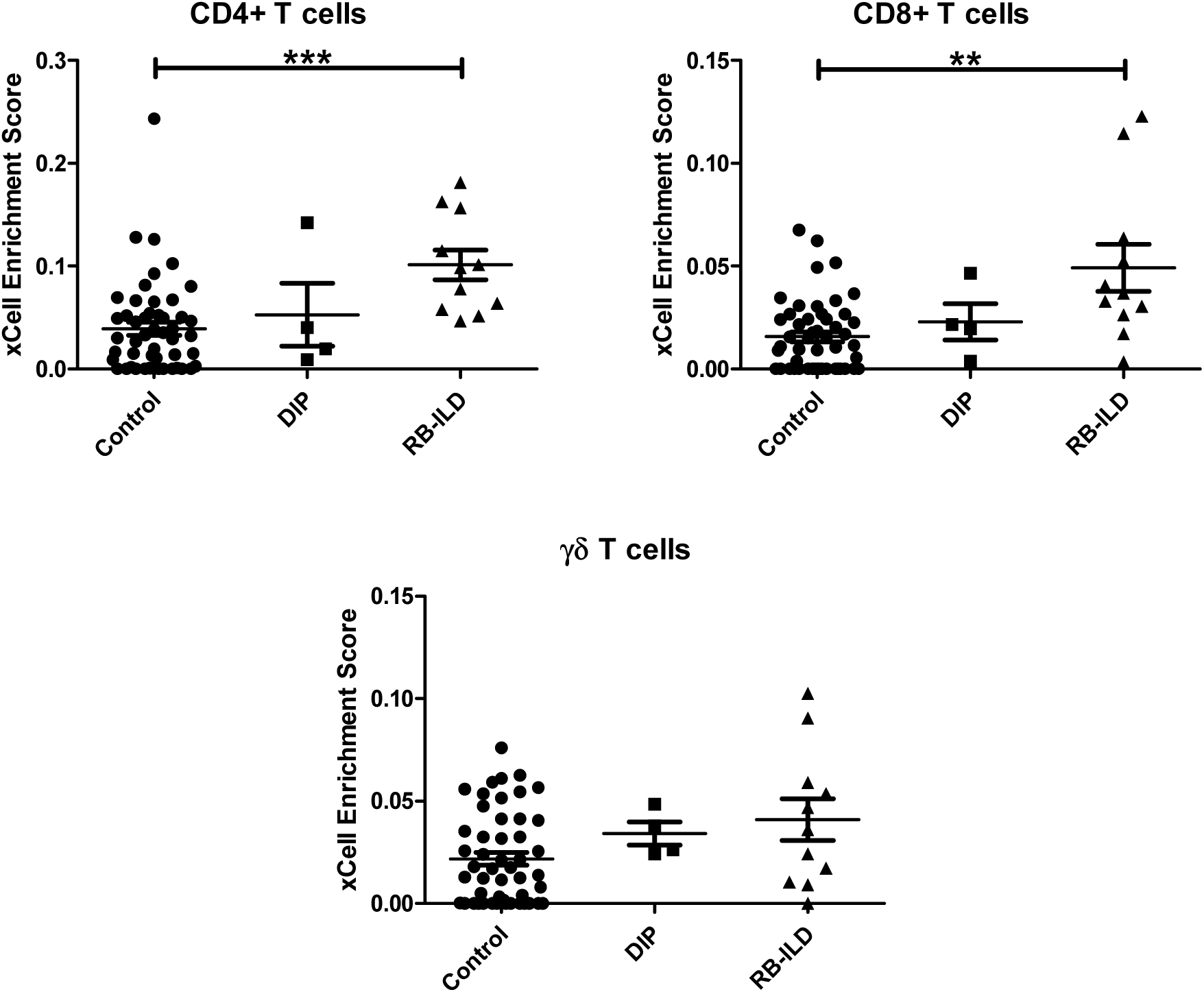
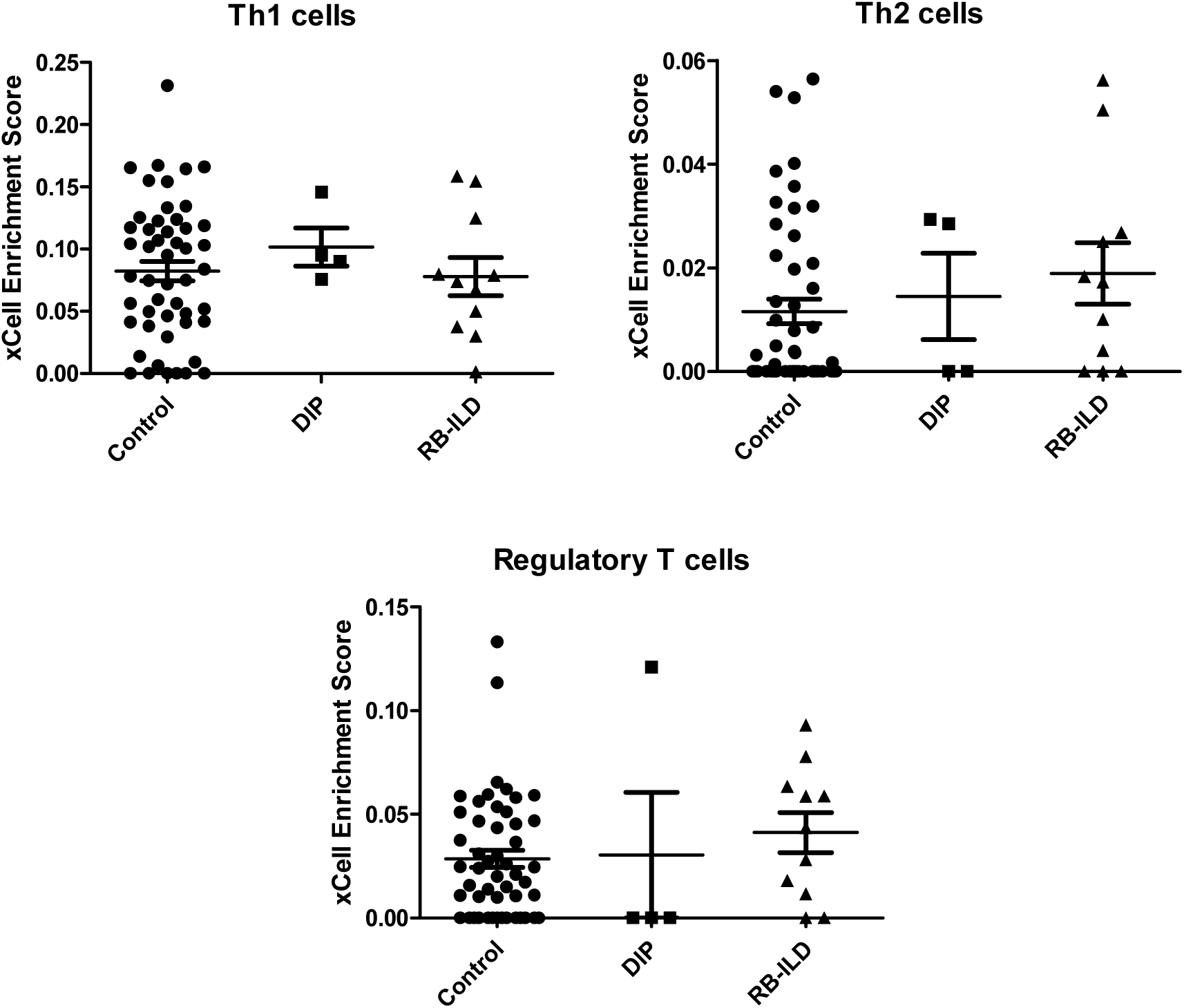
T cells.

**Figure 6.**
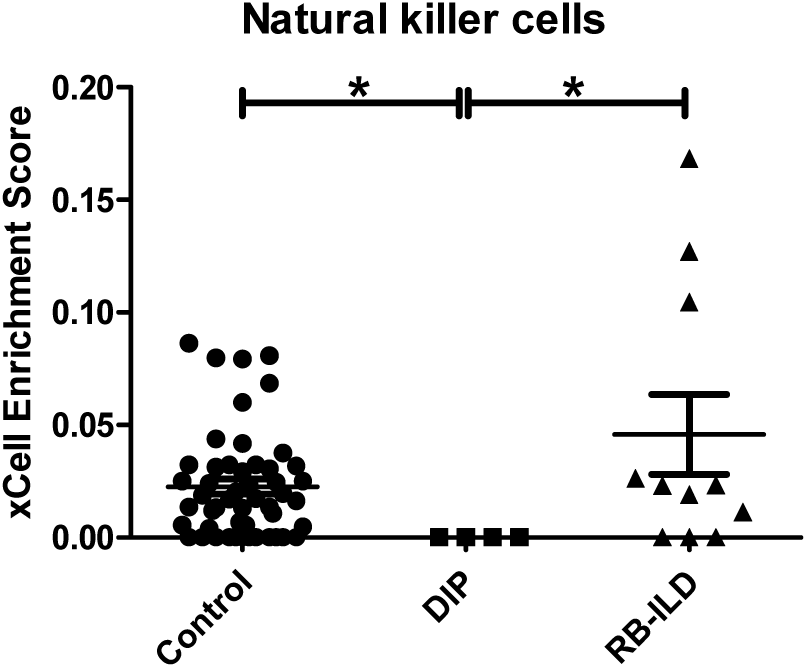
NK cells.

#### Myeloid cells

The gene signature of macrophages (both M1 and M2) was significantly enriched in DIP compared to control (Figure 7). A gene signature of dendritic cells was also significantly enriched in DIP and RB-ILD compared to control. This appears to be driven by conventional dendritic cells rather than plasmacytoid dendritic cells (Figure 8). Although IPA predicted that the “Dendritic Cell Maturation” pathway is strongly activated in DIP compared to control (Table 2), xCell suggested that the gene signature of immature dendritic cells, rather than activated dendritic cells, was significantly enriched in DIP (Figure 8).

**Figure 7.**
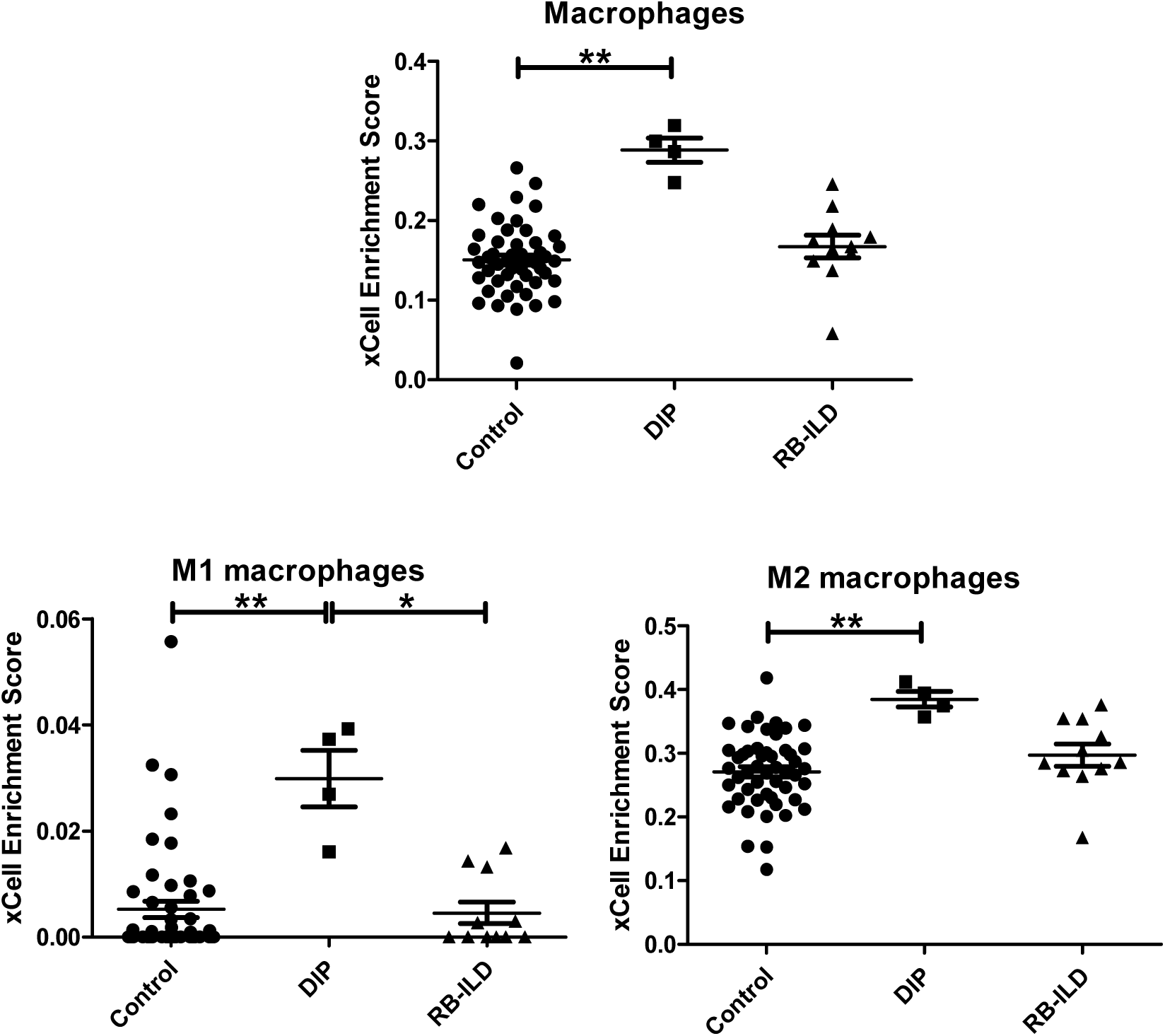
Macrophages.

**Figure 8.**
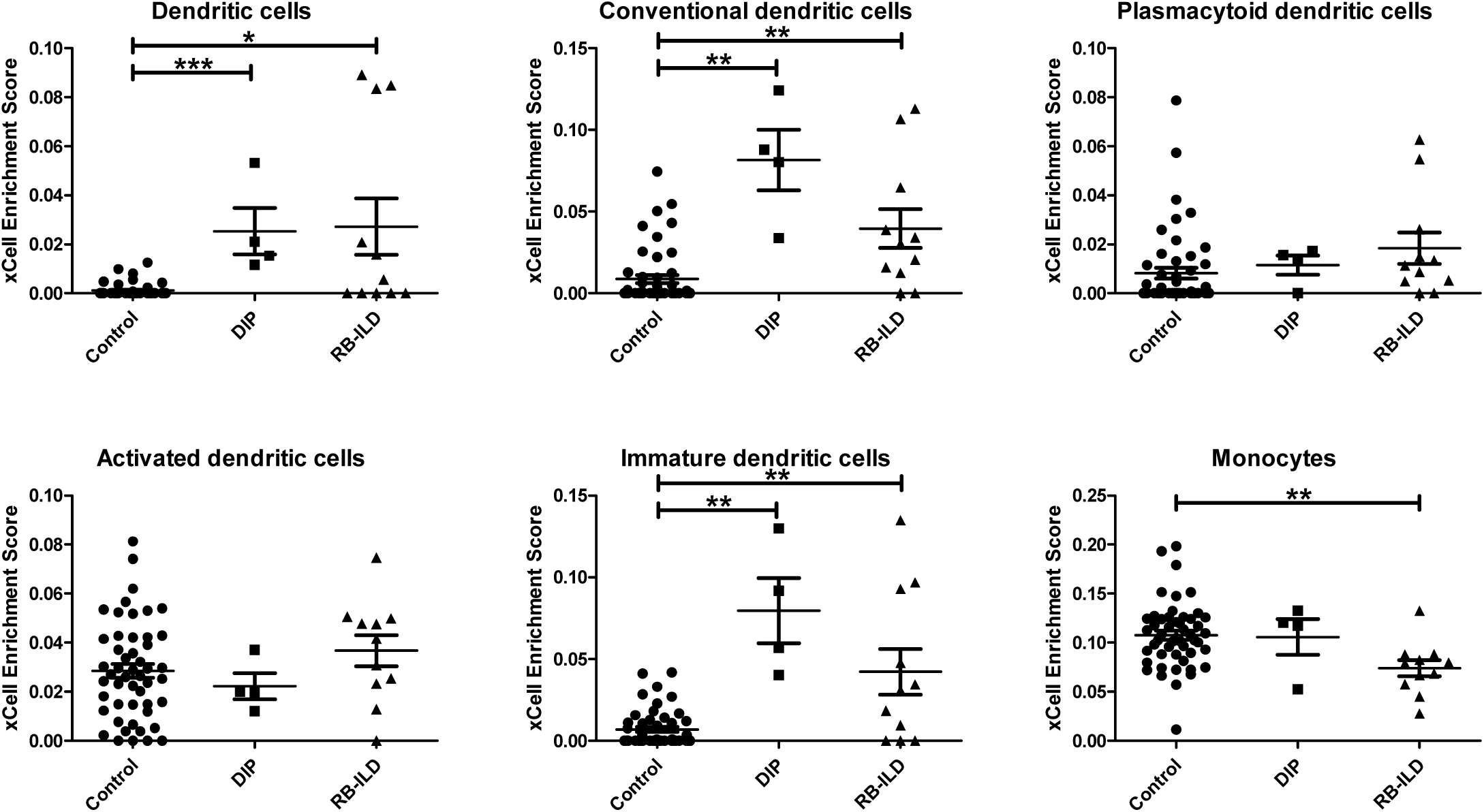
Dendritic cells.

The gene signature of neutrophils was significantly less enriched in DIP and RB-ILD compared to control (Figure 9). In contrast, the gene signature of eosinophils was significantly enriched in DIP compared to RB-ILD and control (Figure 9). This is consistent with previous studies reporting increased eosinophils in DIP (11).

**Figure 9.**
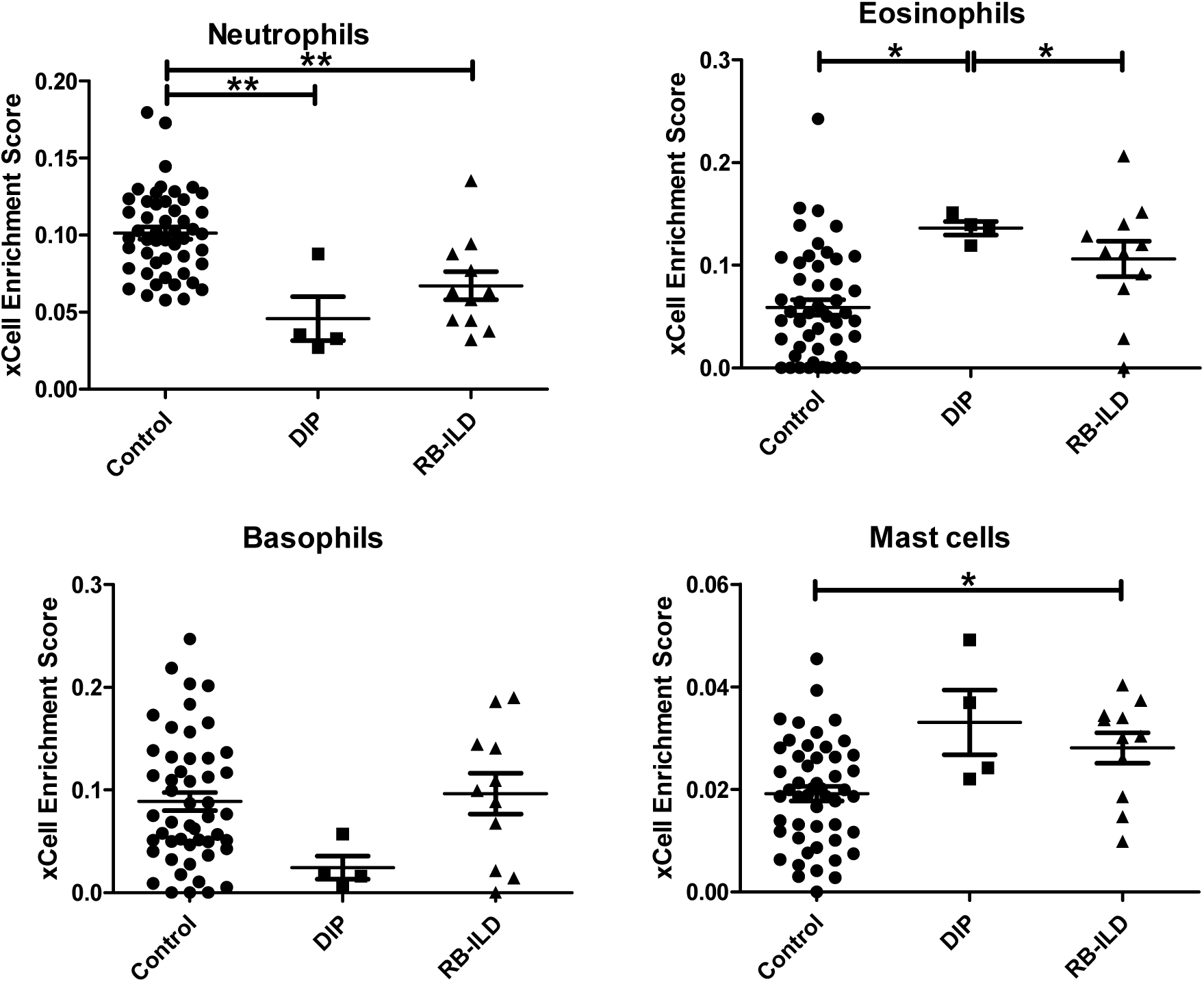
Neutrophils, eosinophils, basophils, and mast cells.

The gene signature of basophils appeared less enriched in DIP compared to RB-ILD and control, but there is no statistically significant difference in the enrichment scores (Figure 9). The gene signature of mast cells appeared more enriched in DIP compared to RB-ILD and control, but there is no statistically significant difference in the enrichment scores (Figure 10). The gene signature of mast cells was significantly enriched in RB-ILD compared to control (Figure 9).

**Figure 10.**
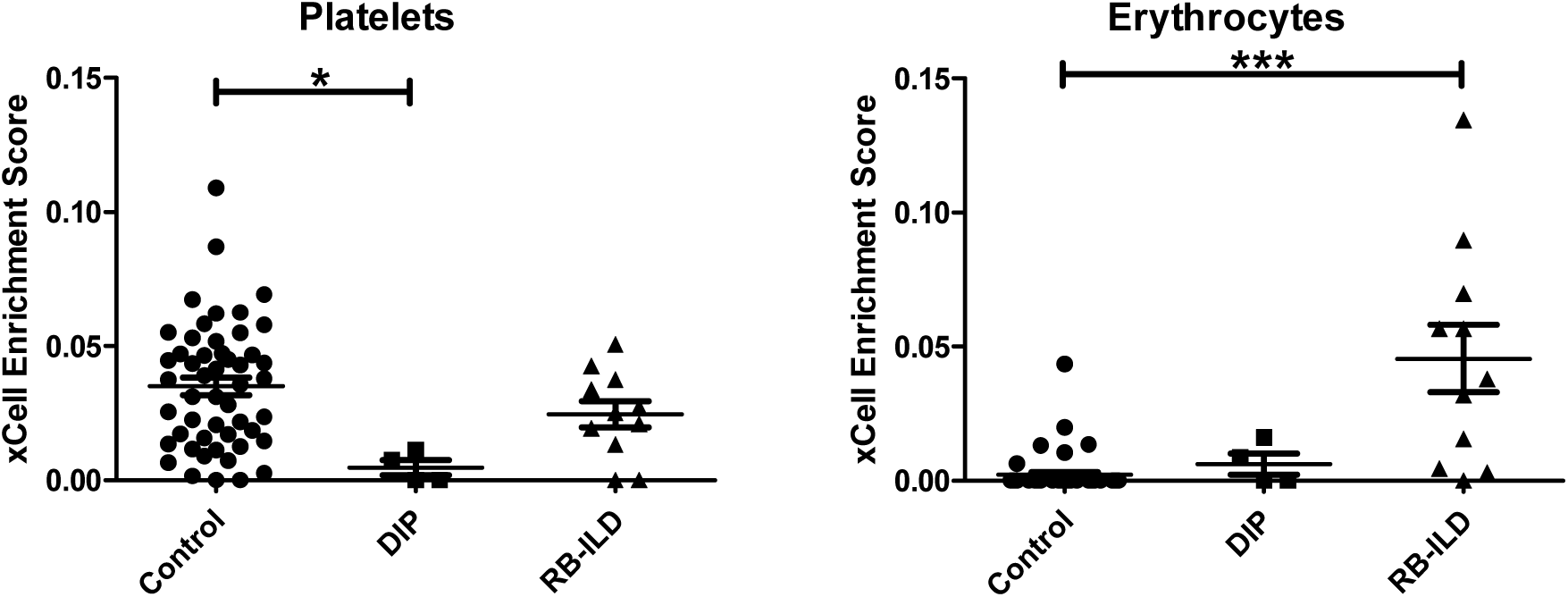
Platelets, and erythrocytes. Please note that xCell scores predict relative enrichment for cell types, not the proportions. Kruskal-Wallis test followed by Dunns multiple comparison tests (all-group comparison). *p<0.05, **p<0.005, ***p<0.001

Interestingly, the gene signature of platelets was significantly less enriched in DIP compared to RB-ILD and control (Figure 10). The gene signature of erythrocytes was significantly enriched in RB-ILD, but not in DIP, compared to control (Figure 10).

## DISCUSSION

Our analysis demonstrates that *CSF2* (encoding GM-CSF) is a critical upstream regulator of the DIP transcriptome. This is in line with the recent finding that lungs of SPC-CSF2 mice mimic DIP (1), and supports the concept that abnormally increased pulmonary GM-CSF signaling may play a critical role in the pathogenesis of DIP.

Our analysis with IPA suggested that many other molecules (e.g., cytokines, transcription factors, and transmembrane receptors) may also be involved in the pathogenesis of DIP. For example, IPA predicted that *SPP1* (encoding osteopontin, a glycoprotein with cytokine-like properties) is another upstream regulator strongly activated in DIP (compared to RB-ILD and control). *SPP1* expression was also significantly increased in DIP compared to RB-ILD and control. Osteopontin has been implicated in the pathogenesis of various lung diseases such as COPD, IPF, DIP, and pulmonary Langerhans cell histiocytosis (PLCH) (9, 12-14). Of note, in both DIP and PLCH, osteopontin concentration is increased in BAL fluid, and BAL cells spontaneously produce high amounts of osteopontin. Furthermore, combined stimulation with osteopontin and GM-CSF enhanced the survival of AMs derived from non-smokers (9). It is quite possible that GM-CSF and osteopontin synergistically contribute to the pathogenesis of DIP in humans. Interestingly, in rodents, pulmonary GM-CSF overexpression caused DIP-like disease (1), whereas pulmonary osteopontin overexpression caused PLCH-like disease (9).

Our analyses with IPA and xCell suggested that immune cells other than alveolar macrophages, both in innate and adaptive immunity, may be involved in the pathogenesis of DIP. For example, the analysis with IPA suggested that the “Dendritic Cell Maturation” pathway is activated in DIP, and the analysis with xCell suggested that the gene signature of dendritic cells is enriched in DIP. This is consistent with previous reports that cigarette smoke induces cytokines such as GM-CSF and osteopontin which can stimulate the recruitment of Langerhans cells (9, 15). It has been shown that GM-CSF has a homeostatic local role in normal tissue macrophage and dendritic cell survival and/or function in the lung (16).

Regarding lymphocytes, the analysis with IPA suggested that the “B Cell Development” pathway is overrepresented in DIP lung. It also predicted that BCR (B cell receptor) and *CD40* (a gene expressed by antigen-presenting cells including B cells) are activated upstream regulators. This appears consistent with previous reports showing that a subset of B cells express GM-CSF receptors and that autocrine production of GM-CSF may contribute to their survival (17, 18). The gene signature of T cells is not significantly enriched in DIP compared to control. However, this does not exclude the possibility that T cells play an important role in the pathogenesis of DIP, especially given that IPA did suggest overrepresentation of many pathways related to T cells in DIP. The gene signature of NK cells is significantly less enriched in DIP compared to RB-ILD and control. This change may be simply due to the effects of cigarette smoke, which has been shown to impair function of NK cells (19). However, this change may also be attributed to altered functions of other types of cells interacting with NK cells. Impaired NK cells may be contributing to the pathogenesis directly or indirectly (by activating other immune cells such as macrophages, B cells, and T cells).

Another interesting finding was that the gene signature for platelets is less enriched in DIP lungs compared to controls. The reason for this is unclear, but may be because platelets in DIP lungs have a very different gene expression profile compared to control lungs. One possibility is that the altered lung environment in DIP affects the platelet transcriptome, as seen in “tumor-educated platelets” in patients with lung cancer (20). The second possibility is that platelet biogenesis is deregulated in DIP lung. Recently, the lung has been shown to be a site of platelet biogenesis and a reservoir for hematopoietic progenitors, including megakaryocyte progenitors (21). The third possibility is that some unknown extrapulmonary factors cause alteration in the DIP platelet transcriptome. This is an interesting question and deserves further research. Another interesting question which merits further research is whether platelets (with altered transcriptome) in DIP actively contribute to inflammatory signaling and subsequent DIP pathogenesis, as seen in other lung conditions (22).

Some drugs and chemicals were predicted by IPA to be upstream regulators. For example, glucocorticoids were predicted to be significantly “inhibited” in DIP compared to control. Again, this may be reflecting the fact that corticosteroids are an effective treatment in some DIP cases. Chloroquine, 4-hydroxytamoxifen, and proteasome inhibitors were also predicted to be significantly “inhibited” in DIP, so we could speculate that these drugs may be effective treatments
for DIP. In fact, it has been reported that a child with DIP due to secondhand cigarette exposure was successfully treated with steroids and hydroxychloroquine (23). In contrast, drugs predicted to be “activated” in DIP such as estrogens and PPAR agonists might be detrimental to DIP. However, it should be noted that these speculations are not based on solid evidence and thus remain in question. It is unclear why filgrastim (granulocyte-colony stimulating factor [G-CSF]) was predicted to be significantly “inhibited” in DIP, despite the fact that G-CSF and GM-CSF have many similarities.

There are several limitations in our study. One of the major limitations is that the sample size for DIP is small (only four cases). Another major limitation is that we were unable to provide validation of microarray analyses (e.g., qRT-PCR for differentially expressed transcripts identified in the microarray analysis) due to a lack of the access to the samples. We also acknowledge that in silico prediction/estimation of gene signature enrichment for various types of cells in tissues is still far from perfect, and is especially difficult in rare types of cells. It remains to be seen if other in silico methods/algorithms outperform xCell, which integrates the advantages of gene enrichment with deconvolution approaches. xCell has been shown to outperform CIBERSORT, a major deconvolution-based method (8, 24), but has not been compared with immunoStates, another deconvolution-based method which has been shown to outperform CIBERSORT (25). At any rate, no in silico algorithm is expected to be perfect in predicting cell composition/enrichment in tissue. Discoveries made using digital dissection methods must be rigorously validated using other technologies (e.g., flow-cytometry, single cell RNA-seq) to avoid hasty conclusions. For example, single-cell RNA-seq may be able to predict cell composition in tissue more accurately. However, it has its own limitations, such as requirement of tissue dissociation, changes in gene expression profile associated with tissue dissociation, and bias towards types of cells that are easily dissociated from tissue (8).

In summary, our analysis revealed that the transcriptome of DIP lungs is distinct from that of RB-ILD and controls. It also supports the notion that abnormally increased pulmonary GM-CSF signaling may play a critical role in the pathogenesis of DIP. Furthermore, it suggested that immune cells other than alveolar macrophages, such as B cells, may be involved in the pathogenesis of DIP.

## Figure Legends

**Supplemental Figure 1.**
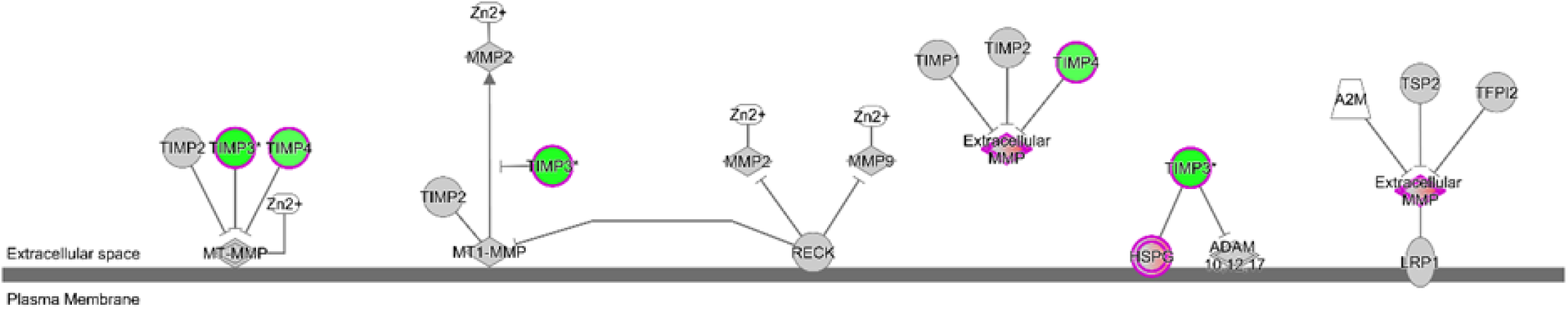
“Inhibition of Matrix Metalloproteases (MMPs)” is predicted to be inhibited in DIP. (i.e., MMPs are predicted to be relatively activated in DIP.)

**Supplemental Figure 2.**
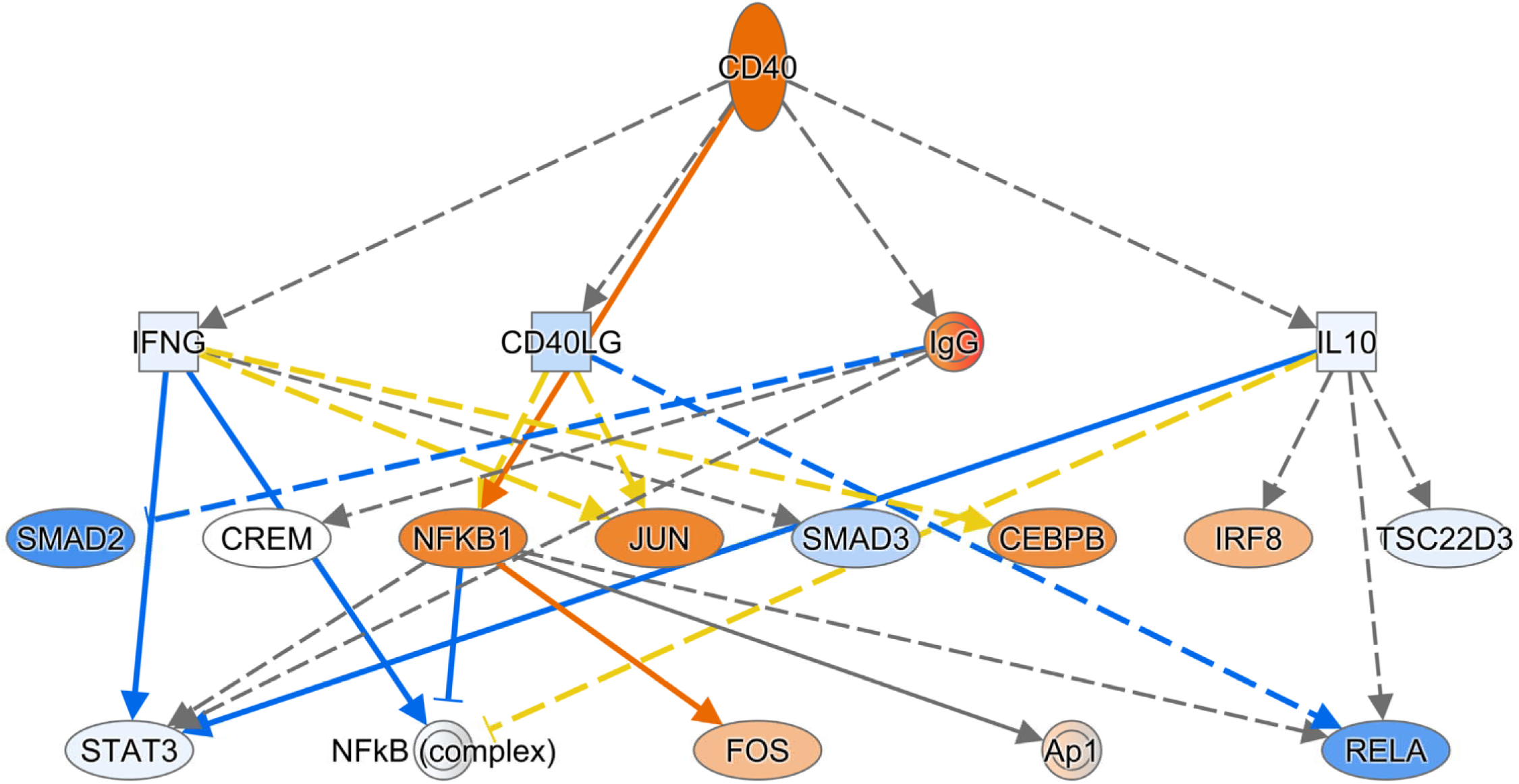
*CD40* is predicted to be one of the activated upstream regulators of the DIP transcriptome.

**Supplemental Figure 3.**
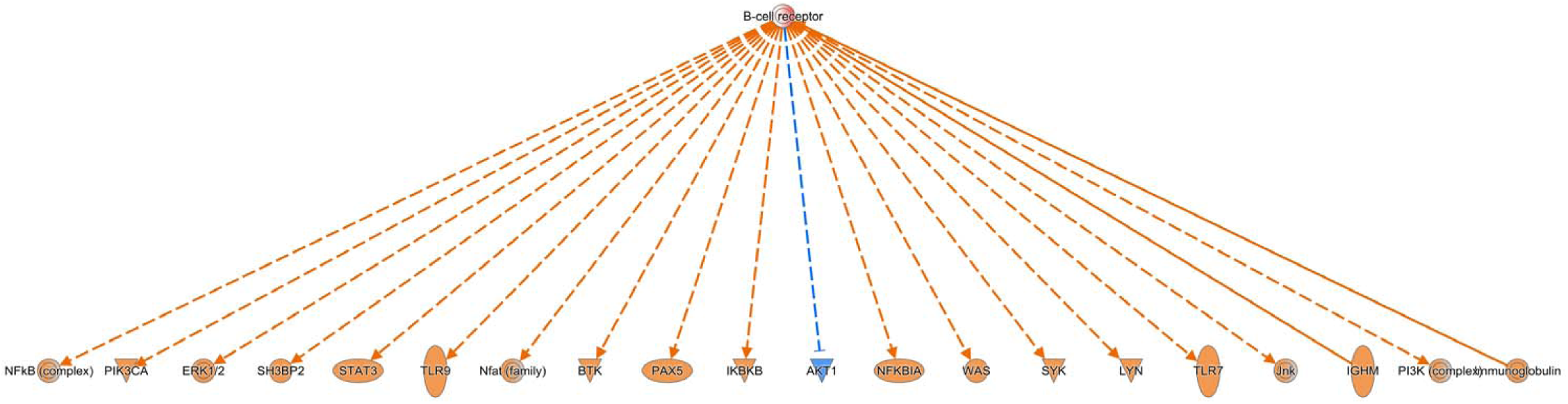
B cell receptor (BCR) is predicted to be one of the activated upstream regulators of the DIP transcriptome.

**Supplemental Figure 4.**
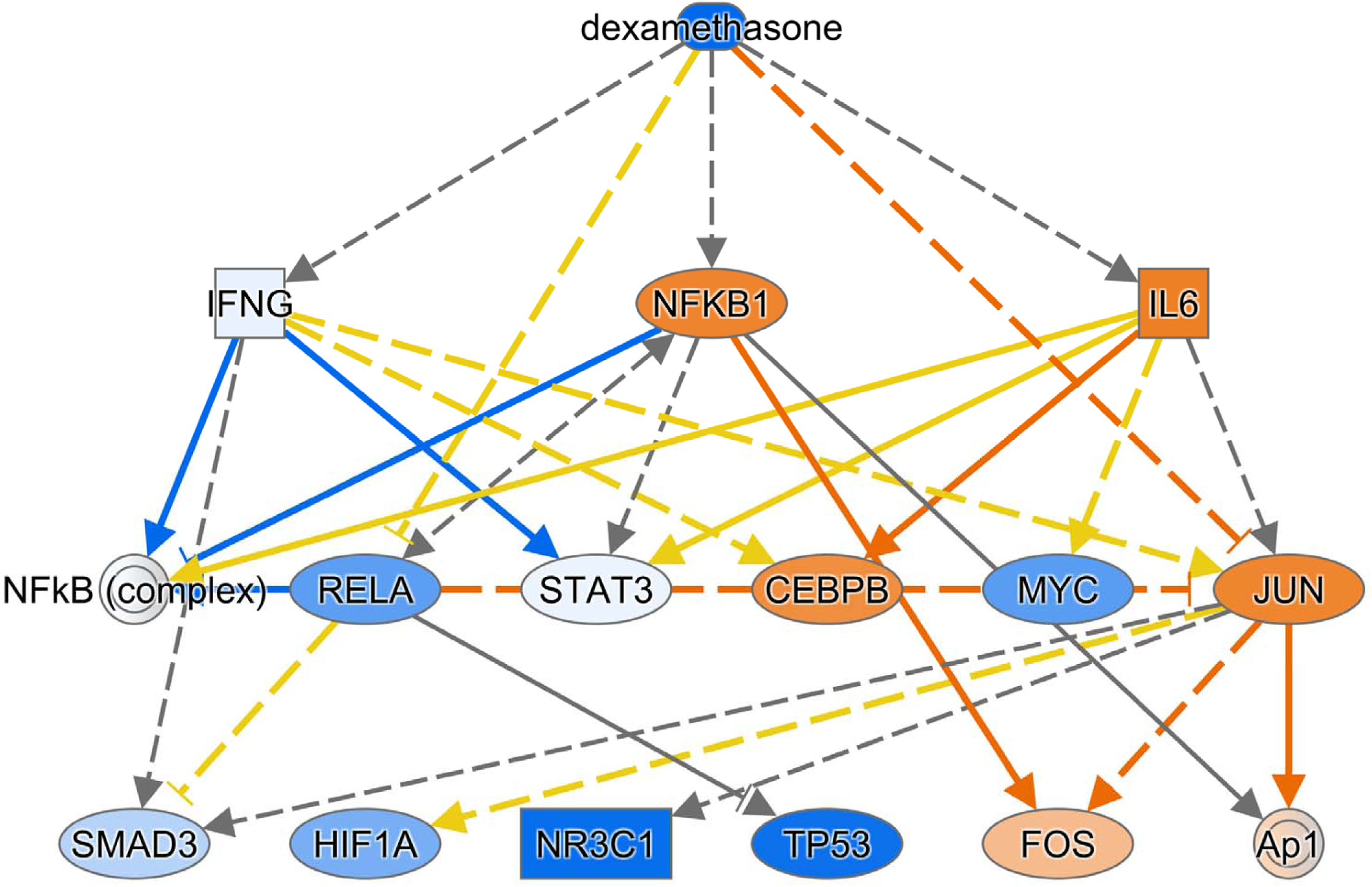
Dexamethasone is predicted to be one of the “inhibited” upstream regulators of the DIP transcriptome.

**Supplemental Figure 5.**
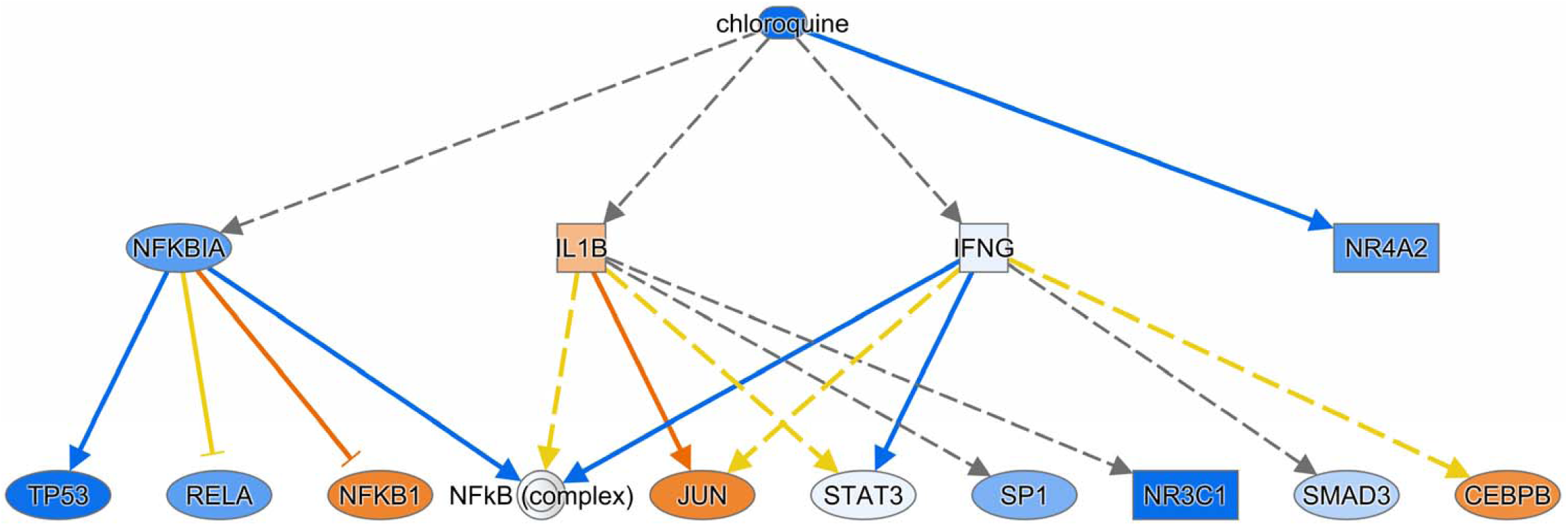
Chloroquine is predicted to be one of the “inhibited” upstream regulators of the DIP transcriptome.

